# Conserved hormonal and molecular mechanisms underlying behavioural maturation in open- and cavity-nesting honey bees

**DOI:** 10.1101/2021.03.25.436783

**Authors:** Sruthi Unnikrishnan, Aridni Shah, Deepika Bais, Ashwin Suryanarayanan, Axel Brockmann

## Abstract

Division of labour in honey bees is based on a process of behavioural development where the worker bee successively performs different tasks at different ages. Workers start with tasks related to brood care and nest maintenance and move on to become foragers. This process of worker behavioural maturation is well studied in *Apis mellifera*. Juvenile hormone is one of the major drivers of this behavioural maturation, which is also accompanied by changes in brain physiology and anatomy including changes in neuronal gene expression and connections of neurons. Recent studies have identified whole networks of genes associated with specific tasks like nursing, guarding and foraging. Based on this detailed knowledge in *A. mellifera*, we ask whether major characteristics of the behavioural maturation process and the underlying hormonal and molecular changes are similar or different in two other honey bee species, the phylogenetically ancestral open-nesting *A. florea* and the more derived cavity-nesting *A. cerana*. Our behavioural studies show that workers of *A. florea* exhibit a slower pace of behavioural maturation and on average start foraging at a later age. However, the basic hormonal and molecular changes associated with onset of foraging are similar between both species. Based on our findings, we propose that evolution of accelerated behavioural maturation in cavity-nesting species is likely attributed to changes in the temporal dynamics of juvenile hormone titres.

## Introduction

Division of labour among individuals is a hallmark of eusocial insects and is believed to be a major factor in their evolutionary success and ecological dominance [1]. Division of labour could be based on physical castes with individuals performing tasks based on their morphological phenotypes or temporal castes with individuals performing tasks based on their age with each individual performing different tasks as they grow older [2–5]. This age-based division of labour or age-polyethism can be considered as a form of behavioural development, similar to juvenile-adult maturation in vertebrates [6], where hormones regulate a succession of behavioural changes by means of changing the structure and physiology of the body and the brain [7].

The Western honey bee, *Apis mellifera*, has been one of the most successful and well-studied social insect species for understanding the behavioural mechanisms as well as the neural and molecular processes underlying worker age-polyethism and behavioural maturation [8,9]. Workers of *A. mellifera* live for about 6 weeks during the summer season. The first half of their lives are spent inside the hive and the second half, outside as foragers. During the first few days after eclosion, workers mostly perform cell cleaning activities in the brood area. After this, the task repertoire increases to include feeding and taking care of brood and nestmates as well as maintaining the nest. In the second and third weeks workers get involved in storing and processing of food, dropping some early life tasks such as cleaning and brood care. Around the end of the third week, workers shift to foraging for nectar and pollen and generally do not perform any other inside-nest tasks till they die [3,10]. Hence, the most striking change in the workers’ life is their transition from doing tasks within the colony to foraging, and this onset of foraging has often been used as an experimental paradigm to study the underlying hormonal and molecular processes involved in age polyethism [11–14].

A key pacemaker of the onset of foraging is juvenile hormone (JH). JH titres in the haemolymph are significantly different between nurse bees and foragers and increase with age [7,15–18]. In fact, treatment with methoprene, a JH analog, accelerates the onset of foraging [16,19–22]. However, allatectomised bees (bees whose corpora allata, which is the production centre of JH, were removed) still developed into foragers, albeit with a delay, indicating that JH is not necessary for the behavioural maturation of honey bee foragers [23]. Developmental processes are generally regulated by several molecular mechanisms working synergistically [24], so there is some experimental evidence that hormones and peptides involved in feeding behaviour, like insulin-like peptide (Ilp) and Neuropeptide Y-like (*NPF*) as well as neuromodulators like octopamine, affect the onset of foraging [25,26]. Thus, it is possible that there is no primary releaser but a network of excitatory and inhibitory molecular pathways that regulate the onset of foraging. An important mutual inhibitory interaction affecting behavioural maturation has been identified between JH and the egg-yolk precursor protein, vitellogenin [12,27–34]. Vitellogenin titres are generally high in nurses and inhibit JH synthesis. However, when JH levels rise, it in turn inhibits vitellogenin synthesis. This results in the increase of JH titres which then accelerates the onset of foraging [34].

In the last fifteen years studies on age-polyethism have focused on studying brain gene expression differences between nurses and foragers as well as changes in gene expression during regular behavioural maturation or during precocious foraging induced by hormone treatment [8,35]. Studies estimate that about 1000 genes are differentially expressed in the brains of nurses and foragers and their expression is regulated by a set of at least 15 transcription factors [36,37]. Some of these identified transcription factors are associated with nursing behaviours, such as broad complex (*BR-C*) and *nautilus* or *MyoD1*, and others with foraging behaviours, e.g. *ultraspiracle* (*usp*), and *egr-1* (early growth-response 1) [13,37–39]. Only for some of these transcription factors there are causal manipulative experiments verifying their function in behavioural maturation.

The detailed knowledge of the hormonal regulation and the molecular changes in the brain associated with worker behavioural development and onset of foraging in *A. mellifera* and the recent sequencing of the genomes of the three major Asian honey bee species, *A. dorsata*, *A. florea* and *A. cerana* [40–42], opens the unique chance to explore variation and evolutionary changes in behavioural maturation and its molecular underpinnings within a small monophyletic group of species [43–45]..

Honey bee species differ in their distribution range, nesting behaviour as well as body and colony size. The phylogenetically ancestral dwarf and giant honey bee species (e.g., *A. florea* and *A. dorsata*), also known as open-nesting honey bees, usually build nests comprising of a single comb attached to tree branches, rooftops or cliff-sides. The workers of the colony form a curtain around the comb, such that it is protected from adverse environmental conditions, parasites and predators [46–49]. Cavity-nesting species such as *A. mellifera* and *A. cerana*, construct their nests inside crevices of rocks or buildings or cavities in tree-trunks, and they usually build multiple combs. Since the cavity protects the nest these species generally do not form a curtain surrounding the combs [46,50,51]. These differences in nesting behaviour are expected to affect the behavioural maturation and age at the onset of foraging of the worker caste. More specifically, it was hypothesised that due to the relatively smaller number of brood cells and the necessity of maintaining a large workforce to sustain the curtain there would be a delay in the transition of individuals from performing nurse tasks to becoming foragers in open-nesting as compared to cavity-nesting honey bees [46,47]. In addition, worker behavioural maturation could be different between temperate and tropical honey bees. Workers of temperate species or populations like *A. mellifera carnica* and *ligustica*, which have been mainly used for the study of age-polyethism, might show an accelerated behavioural development because more foragers are required to build up sufficient food stores to survive the winter phase [52,53].

In the current study we have performed a comparative study of the behavioural maturation between an open nesting species, *Apis florea* and a cavity nesting species, *Apis cerana* both of which are tropical honey bees. Our objectives were 1) to study whether there is a difference between open- and cavity-nesting honey bees and 2) to understand whether tropical species follow the same patterns for behavioural maturation as the temperate species, *A. mellifera*. First, we studied the foraging behaviour to identify the age of onset of foraging for both species. Second, we studied the temporal dynamics of juvenile hormone and vitellogenin in both the species. Third, we studied the juvenile hormone titres, *Vg* and insulin-like peptide *(ilp-1)* levels in nurses and foragers. Finally, we studied the expression levels of four transcription factors in nurses and foragers, two of which are associated with foraging behaviour, *usp* and *egr-1* and two of which are associated with nursing behaviour, *BR-C* and *nautilus*.

## Methods

### Behavioural observations

#### Apis cerana

Four-framed colonies of *Apis cerana* were obtained from a local bee keeper and maintained in the campus of the National Centre for Biological Sciences, TIFR, Bangalore. For the behavioural experiments the colonies were split and two frames with the queen were transferred into an observation hive. The brood frame from the remaining half of the colonies were kept inside incubators at a temperature of 33-34°C for 24 hours. This allowed late-stage pupae to eclose as adults. These day-olds were then individually colour marked and released into their respective colonies. Only two colonies could be successfully studied. Five colonies absconded before end of the experiment and could not be observed completely. Out of the two, for the 1^st^ colony (Cerana 1) 77 marked day-olds and for the 2^nd^ colony, Cerana 2, 164 marked day-olds were released into the colonies. Behavioural observations were done to follow the behaviour of these marked bees across days till day 24. A video camera kept at the entrance of the observation hive for 2-4 hours every day, till day 35 allowed to study when the introduced bees started foraging. An individual was considered to be a forager if their foraging trip duration was equal to or more than 1.75 minutes. This flight duration equals the least amount of time taken by a pollen forager. In addition, bees were also considered as foragers if they followed a dance or danced. In this manner the age of first foraging trip or onset of foraging of each marked bee was identified. The first colony, Cerana 1 was observed between December 2017 – February 2018 and the second colony, Cerana 2 between April-June 2019.

#### Apis florea

Colonies of *Apis florea*, were collected with the help of a local bee keeper and maintained on the campus of the National Centre for Biological Sciences, TIFR, Bangalore. Brood combs from separate colonies (not the colonies used for the study) were kept in the incubator at a temperature of 33-34°C for 24 hours for adult bees to eclose. These day-old bees were individually colour marked (100 bees for each colony) and introduced into the colonies. A video camera kept in front of the colonies for 2-4 hours every day till the colony absconded, recorded the flight activity of the individuals and enabled us to study the foraging behaviour of the marked bees. Similar to *A. cerana*, foraging trips were studied and the onset of foraging of the marked bees was identified. Out of 10 colonies, only two (Florea 1 and Florea 2) were successfully observed as the remaining absconded. The first colony of *A. florea*, Florea 1, absconded by day 70 and Florea 2 by day 40. Florea 1 was observed between December 2018 – February 2019 and Florea 2 was observed between October – November 2018.

### Hormonal and genetic analysis

#### Sample collection

##### (i) Age-related changes in JH titres and *Vg* expression

Day-old bees were marked and released (100 each) in the same manner as for behavioural observations, for 2 colonies each of *A. cerana* (6 frames each) and *A. florea*. The only difference was that *A. cerana* colonies were not transferred into observation hives as in the case of the behavioural analysis. Similar to the behaviour observations, foraging behaviour was studied with the help of video cameras kept outside the colonies for 2-4 hours every day till the end of the collection. Individuals were collected on 0 (day-old bees), 5, 10, 18, 25, 30, 35, 40, 45 and 50 days for measuring their juvenile hormone titres in the haemolymph. After haemolymph extraction these bees were stored in −80°C until measuring abdominal gene expression levels of vitellogenin. The collections and observation of these colonies were done simultaneously between 26^th^ November 2019 to 15^th^ January 2020. All 4 colonies used for JH analysis were different than the ones used for behavioural observations in the previous section. A list of colonies used for the different studies is provided in Table S1.

##### (ii) Nurse-Forager comparison

Individual nurse bees and foragers were collected from 6 colonies each of *A. cerana* (5 frames each) and *A. florea* based on the tasks they were observed to be performing irrespective of their age. Nurses were identified as those bees that inserted their heads into cells containing larvae [54–57]. Foragers were identified based on the fact that they had pollen load when returning to the colony. Three colonies were used for measuring JH titres and the remaining three for measuring *Vg*, *ilp-1* and TF gene expression levels. Individuals collected (except those used for JH studies) were immediately stored at −80°C.

#### Juvenile Hormone measurements

##### (i) Extraction of haemolymph and sample preparation

The collected individuals were immediately anaesthetized on ice. Their antennae were cut using dissection scissors and the individuals were inverted and centrifuged, such that their haemolymph would flow out into collection tubes through the cut antennae [58]. 10μl of this collected haemolymph from each individual bee was then mixed with 22μl methanol and 8μl of internal standard (JHEE). JHEE was made by a transesterification process of JH following methods established by Scholl et al. 2014 [59] and was used as the internal standard. If unable to obtain 10μl of haemolymph from the individual bee, the possible amount that could be collected was taken and then methanol was added up to 10μl, so that the total volume of the mixture came to 40μl. The exact amount of haemolymph added was noted for normalising the differences between bees in this aspect. These samples were then sonicated for 2 minutes and centrifuged for 10 minutes at 14,000 rpm. The supernatant was then analysed using LC/MS-MS.

##### (ii) LC/MS-MS conditions

A Thermo Scientific TSQ Vantage triple stage quadrupole mass spectrometer (Thermo Fisher Scientific, San Jose, CA, USA), connected to an Agilent 1290 infinity series UHPLC system (Agilent Technologies India Pvt. Ltd., India) was used for the LC/MS-MS. The column oven was set at 40°C and the autosampler tray at 4°C. The mobile phase solvent A was 10mM Ammonium acetate with 0.1% Formic acid and mobile phase solvent B was Methanol (100%). The chromatography was carried out in an 80Å column (30 mm × 4.6 mm i.d.), which was protected by a C18 guard column (4mm × 2mm i.d.), both from Phenomenex [58]. Gradient elution was performed at a flow rate of 0.4 ml/min at a column temperature of 40°C from 65 to 100% B within 7 minutes, followed by 100% B for 1.10 minutes and reconditioning at 65% for 3 minutes. The injection volume was set as 10μl.

An ESI-MS/MS was performed using multiple reaction monitoring (MRM) and the reactions 267 to 147 for JH and 281 to 189 for JHEE were followed. The ESI source was operated in positive electrospray mode (ESI+) at 60°C, at spray voltage of 3500 V. Argon was used as the collision gas. The capillary temperature was set at 300°C, sheath gas pressure at 15 and the auxiliary gas at 15. The S-lens RF amplitude for ion 267 was 50.93 and for ion 281 was 47.76. The collision energy was 13 V for the reaction of ion 267 to yield the product ion 147 and 12 V for the reaction of ion 281 to yield the product ion 189. JH eluted out at 4.9 minutes and JHEE at 5.3 minutes, respectively. The calculated amount obtained from MS measurements were multiplied by 4 (as 10 μl out of 40 μl was injected into the LC-MS/MS) and divided by the amount of haemolymph collected for each individual during the statistical analysis.

#### Gene expression measurements of Vitellogenin (Vg), Insulin-like peptides (ilp-1) and transcription factors (usp, Egr-1, BR-C and nautilus)

##### (i) Brain dissection, RNA isolation, cDNA preparation and quantitative PCR

Honey bee brains were first dissected on a dry ice platform in a glass cavity block filled with 100% ethanol. The dissected brain and the corresponding abdomen of the bee were collected in individual tubes. The brain and the abdomen were then homogenized in TRIzol (Invitrogen, Life Technologies, Rockville, MD, USA) with a motorized homogenizer, followed by RNA extraction using the Trizol-chloroform method. Glycogen (20mg/ml, Thermo Scientific, Life Technologies, Rockville, MD, USA) was added to increase the recovery of RNA. The extracted RNA was converted to cDNA using the SuperScript™ III First-Strand Synthesis System (Invitrogen, Life Technologies, Rockville, MD, USA). The qPCR was performed following the same protocol as in Singh et al. [60]using Kapa SybrGreen (Kapa Biosystems, Wilmington, Massachusetts, USA). The standard curve method was followed with *AcRP49* and *AfRP49* as the internal control for *A. cerana* and *A. florea* respectively.

##### (ii) Primers

Primers for *usp*, *egr-1*, *Amilp-1*, *Vg* and *RP49* were designed based on *A. mellifera* primers used in previous literature (Table 1). The *A. mellifera* primer sequence was aligned on the respective *A. cerana* and *A. florea* genes using NCBI BLAST. After aligning the primers, differences in nucleotides, if any were corrected such that the primer sequence matched 100% with the respective gene sequence. No literature source was found for *AmNau* and *A. mellifera* derived *BR-C* primers showed poor efficiency and gave multiple peaks. Hence, new primers were designed for *nautilus* and *BR-C*. The primers for the different genes are given in Table 1.

**Table 1:**
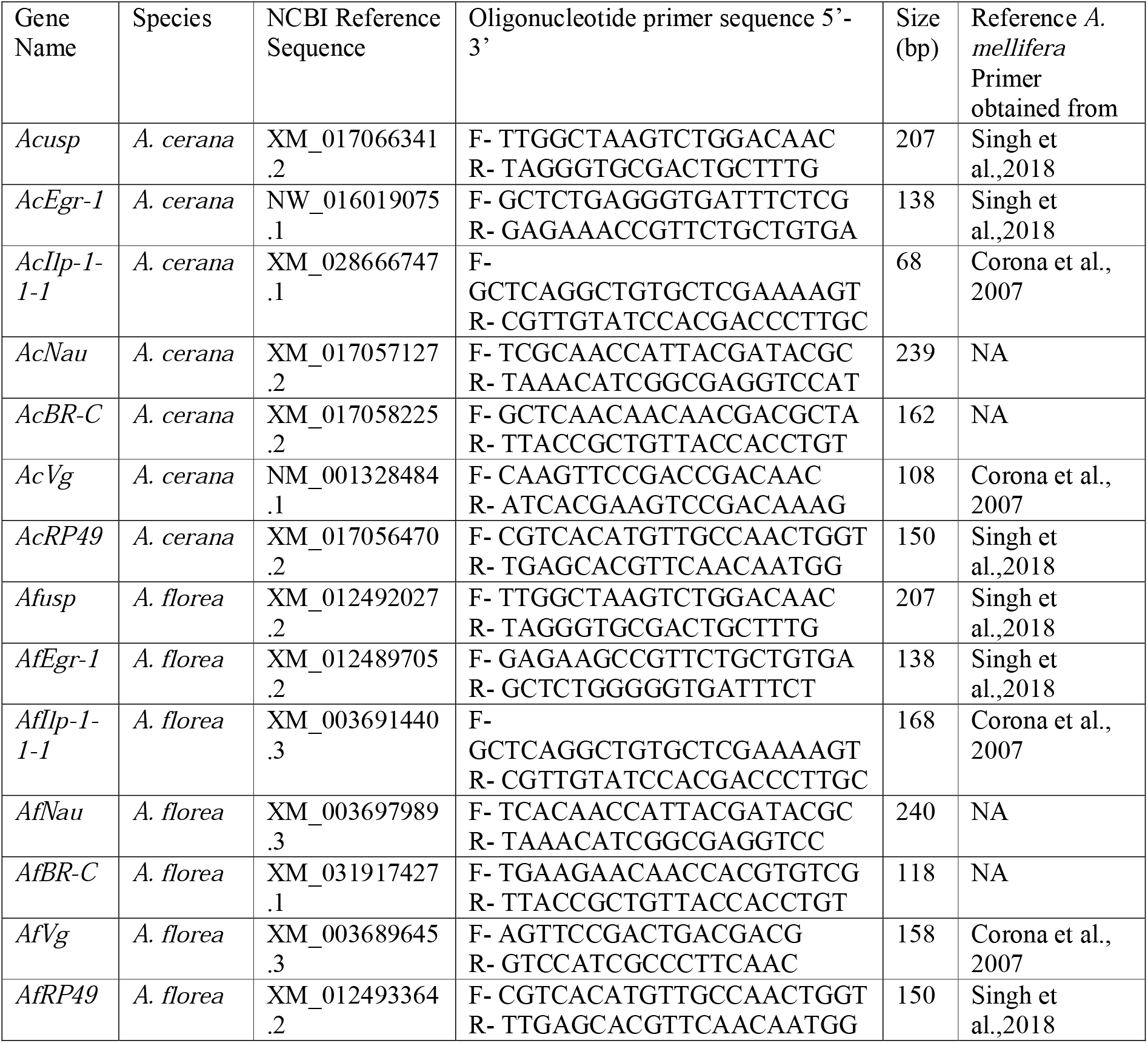
Primers for the transcription factors, *Vg, Ilp-1-1* and *Rp49*.

### Statistical analysis

All statistical analysis was done in R version 3.4.1 [61] with the R studio IDE [62]. For the onset of foraging, a linear mixed effects model with species (two levels, *A. florea* and *A. cerana*) and an interaction term between species and age as fixed effects and the cumulative percentage of bees that initiated foraging as the response variable was built using the lme4 package [63]in R. The nests were the random effect. For the age-based changes in JH levels, two separate mixed effects models for *A. florea* and *A. cerana* was built using the gamlss package in R [64]. Age and foraging status (whether a bee was a forager or not) were the fixed effects and the JH titres were the response variable with colony and batch number (of using the MS machine) as the random effects fitted with a Weibull distribution. For the analysis of the temporal dynamics of vitellogenin levels, separate mixed effects models were built for *A. florea* and *A. cerana*. In both cases, the fixed effect was age and vitellogenin gene expression levels was the response variable with colony as the random effect. For *A. cerana* a lognormal distribution was fit to the model using the lme4 package and for *A. florea* a Weibull distribution was fit using gamlss package. For the comparisons of JH titres between nurses and foragers, separate mixed effects models were built for *A. florea* (fitted with a normal distribution) and *A. cerana* (fitted with lognormal distribution) with JH titres as the response variable and the behavioural state (nurse and forager) as the fixed effect and colony as a random effect. For the gene expression comparisons (*Vg, ilp-1, usp, egr-1, BR-C* and *nautilus*) between nurses and foragers, the values were scaled to normalise for colony variation and linear regression models were fit using the MASS package [65]. Separate models were built for each species and for each of the response variables, i.e., expression levels of *Vg*, *ilp-1* and the 4 transcription factors, *usp, egr-1, BR-C* and *nautilus*, with the behavioural state, nurse and forager, as a categorical predictor variable. The correlations were analysed using Kendall’s correlation coefficient and ggplot2 [66]was used for data visualization. Dharma package [67] was used to test model assumptions and model comparisons were made using AIC values.

## Results

### Onset of foraging in *A. florea* and *A. cerana*

We found that the onset of foraging is much more variable in *A. florea* as compared to *A. cerana* with higher inter-individual variation in *A. florea* than in *A. cerana*. In fact, the age of the very first initiation of foraging is quite similar in both species. In colony Cerana 1 of *A. cerana*, the very first foraging was initiated at the age of day 10 and by day 35, 62% (48/77) bees had become foragers (Fig. 1); 35% (27/77) bees had disappeared before day 22 (see Fig. S1) and 2.5% (2/77) that were still alive at day 24 and had not become foragers by day 35. In colony Cerana 2 of *A. cerana*, foraging was observed for the first time on day 4 and by day 35, 60% (98/164) had become foragers (Fig. 1); 36.5% (60/164) bees had disappeared before day 22 (see Fig. S2) and 3.6% (6/164) bees were still alive at day 24 but had not become foragers by day 35. In *A. florea* in colony Florea 1, by day 7 the first individual had initiated foraging and it was by day 70 that 50% (50/100) of bees had become foragers. In the case of colony Florea 2 of *A. florea*, by day 4 the first foraging was initiated and by day 40 only 20% (20/100) had become foragers (Fig.1). The slope of cumulative percentage of individuals initiating foraging was steeper for *A. cerana* compared to *A. florea*. (Table 2, Fig. 1). These percentages are out of the total number of marked bees released into each colony. Percentage of bees that became foragers out of the total number of bees remaining in each colony is given in the supplementary (Fig. S3).

### Temporal dynamics of JH and *Vg* expression

JH significantly increased with age in both *A. florea* and *A. cerana*. *Vg* did not change significantly with age for either species. There was also no significant correlation found between JH and *Vg*.

**Table 2:** Table showing the fixed effects, random effects and the estimate with standard error, confidence intervals and P values for the linear mixed effects model studying the onset of foraging behaviour in *A. florea* and *A. cerana*. The interaction term between age (in days) and species was significant for both species.

| Fixed effects | Estimate | Std. Error | CI | P value |
| --- | --- | --- | --- | --- |
| (Intercept) | -6.69 | 3.89 | -16.2 – 2.8 | 0.103 |
| speciesFlorea | 1.47 | 5.04 | -9.89 – 15.99 | 0.775 |
| age: Cerana | 2.102 | 0.15 | 1.8 – 2.4 | < 0.0001**** |
| age: Florea | 0.808 | 0.07 | 0.85 – 1.15 | < 0.0001**** |
| Random effects | Variance |  |  |  |
| Nest | 8.79 |  |  |  |
| Residual | 30.13 |  |  |  |

**Fig 1.**
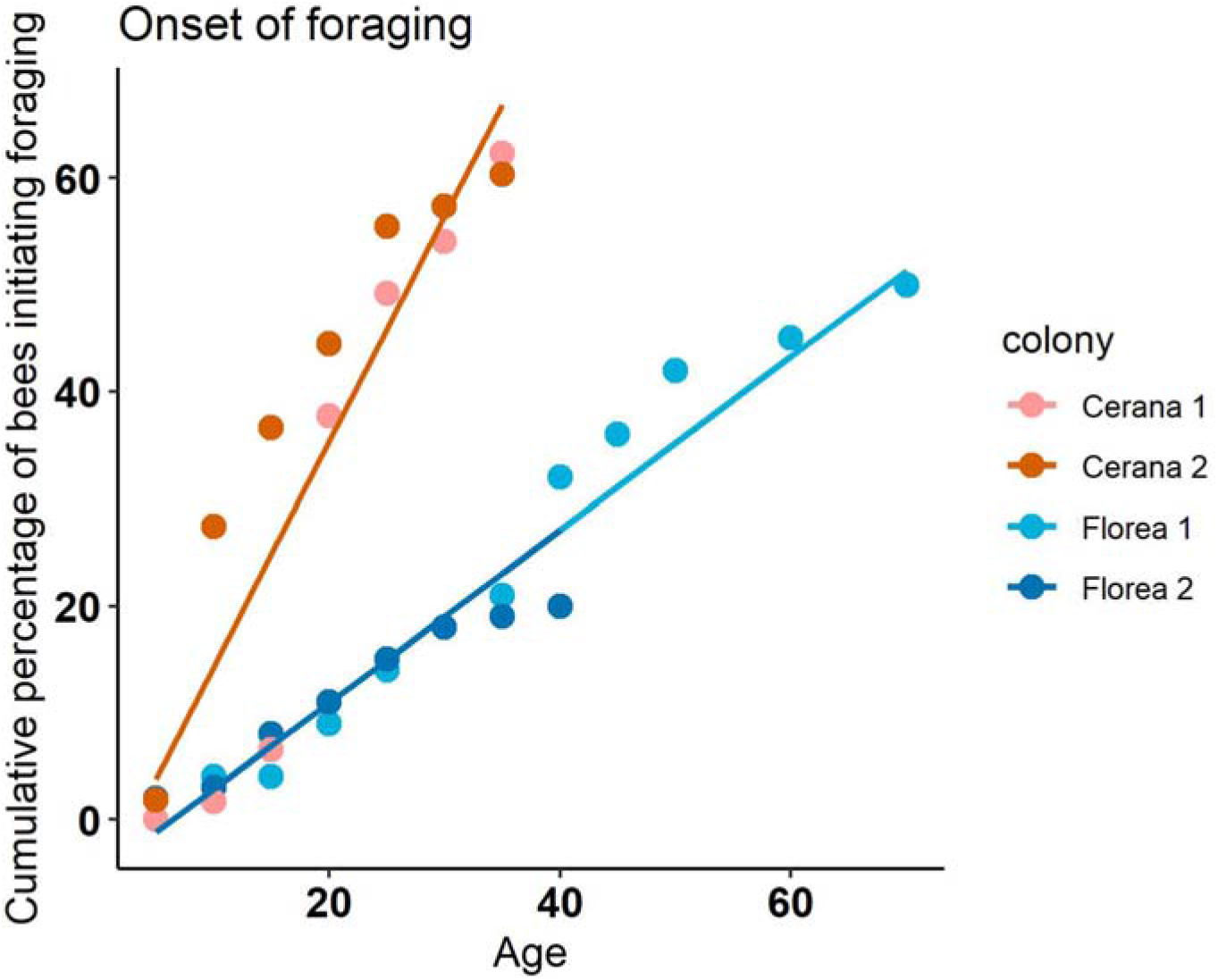
Illustrates the onset of foraging in two colonies each of *A. florea* (shades of blue) and *A. cerana* (shades of red). × axis is the age in days and Y axis indicates the cumulative percentage of bees that initiated foraging calculated out of the total number of marked bees released into the colony. A linear mixed effects model with species and interaction term between age and species as fixed effects, cumulative percentage as response variable and colony as random effect was built. The interaction term was significant for both species, indicating the difference in the speed of foraging initiation between the two species.

#### Juvenile Hormone

In the case of *A. florea and A. cerana*, age as well as foraging status had a significant effect on the JH titres. The interaction term between age and forage status was not significant (Table 3, Fig. 2).

**Table 3:** Table showing the fixed effects, random effects and the estimate with standard error, confidence intervals and P values for the generalised linear mixed effects models (fitted with Weibull distribution) to study the changes in JH titres in haemolymph with age and forage status (whether a bee is a forager or not) for (a) *A. florea* and (b) *A. cerana.* Age and forage status were significant predictors for JH titres in both *A. florea* and *A. cerana*.

| Fixed effects | Estimate | Std. Error | CI | P value |
| --- | --- | --- | --- | --- |
| (Intercept) | -5.52 | 0.153 | -5.82 – -5.22 | <0.0001**** |
| age | 0.055 | 0.0056 | 0.044– 0.066 | <0.0001**** |
| forage.st.F | 3.47 | 1.58 | 0.37 – 6.57 | 0.033* |
| age:forage.st.F | -0.054 | 0.045 | -0.14 – 0.03 | 0.238 |
| Random effects | Std.Dev |  |  |  |
| Nest | 0.22 |  |  |  |
| Batch | 0.56 |  |  |  |
| Residual | 1.02 |  |  |  |

(b) *A. cerana*:
| Fixed effects | Estimate | Std. Error | CI | P value |
| --- | --- | --- | --- | --- |
| (Intercept) | -5.19 | 0.31 | -5.81 – -4.58 | <0.0001**** |
| age | 0.066 | 0.019 | 0.03 – 0.1 | 0.0015** |
| forage.st.F | 2.06 | 0.78 | 0.53 – 3.6 | 0.011* |
| age:forage.st.F | -0.035 | 0.03 | -0.09 – 0.02 | 0.23 |
| Random effects | Std.Dev |  |  |  |
| Nest | 2.81e-05 |  |  |  |
| Batch | 0.0001 |  |  |  |
| Residual | 1.22 |  |  |  |

**Fig 2.**
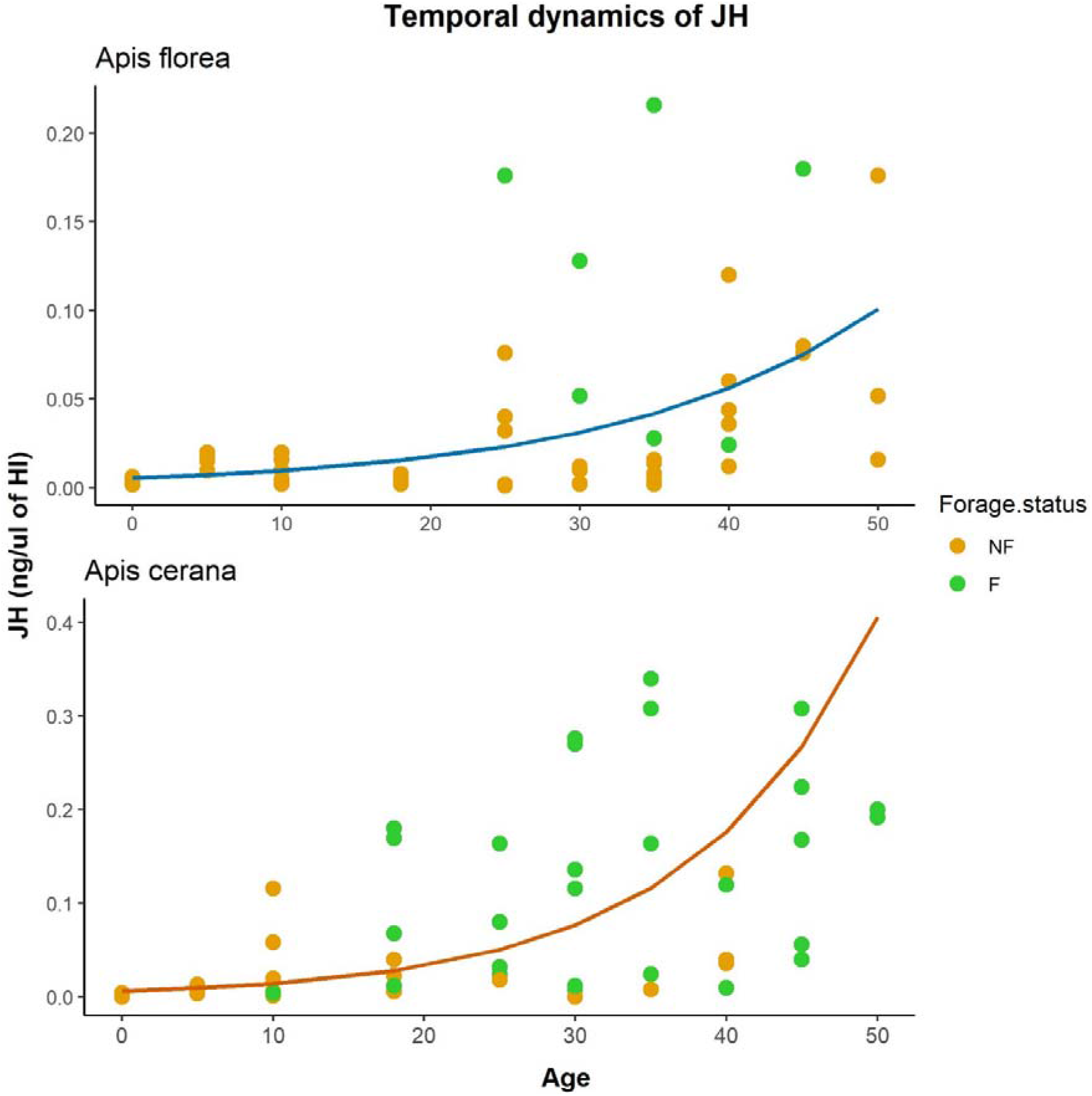
Illustrates the changes in JH titres with age. × axis is the age measured in days and Y axis is the JH titres (in nanogram per microlitre of haemolymph) of the bees. The top panel shows the results for *A. florea* and the bottom for *A. cerana*. Each dot indicates a single bee and orange dots are the bees that had not yet become foragers (NF) by the time of collection while green are the ones that had become (F). Mixed effects models with age and forage status as fixed effects, JH titres as response variable and colony and batch number as random effects were built. Lines indicate the prediction from the models for the relationship between age and JH. In both species, age and forage status had a significant effect on JH levels.

#### Vitellogenin

There was no significant effect of age on *Vg* levels for both *A. florea* and *A. cerana* (Table 4, Fig. 3).

**Table 4:**
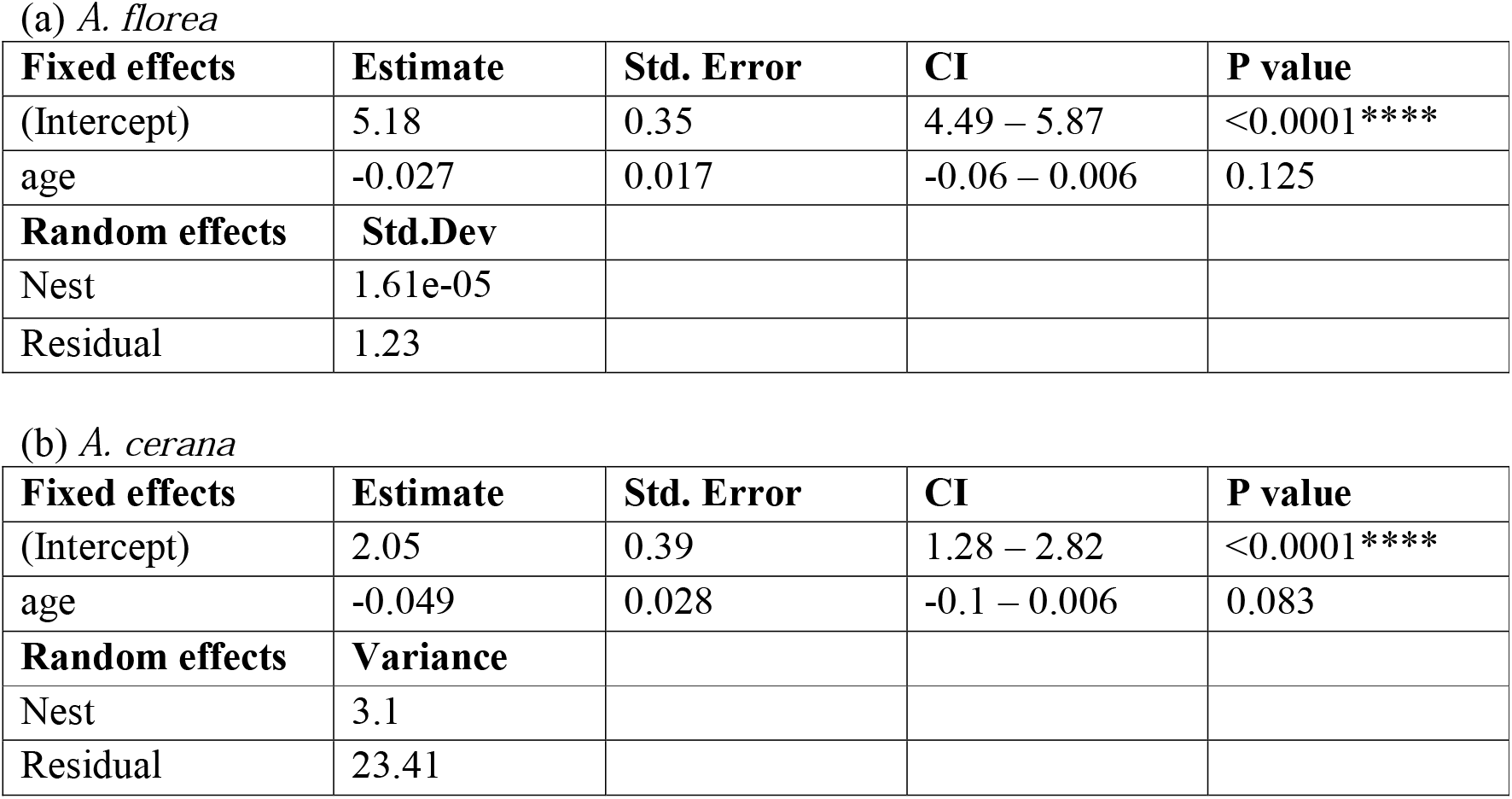
Table showing the fixed effects, random effects and the estimate with standard error, confidence intervals and P values for the generalised linear mixed effects models (fitted with Weibull distribution) to study the changes in *Vg* gene expression levels in abdomen with age for *A. florea* and *A. cerana*. Age was not a significant predictor of *Vg* levels in both *A. florea* and *A. cerana*

| Fixed effects | Estimate | Std. Error | CI | P value |
| --- | --- | --- | --- | --- |
| (Intercept) | 5.18 | 0.35 | 4.49 – 5.87 | <0.0001**** |
| age | -0.027 | 0.017 | -0.06 – 0.006 | 0.125 |
| Random effects | Std.Dev |  |  |  |
| Nest | 1.61e-05 |  |  |  |
| Residual | 1.23 |  |  |  |

(b) *A. cerana*
| Fixed effects | Estimate | Std. Error | CI | P value |
| --- | --- | --- | --- | --- |
| (Intercept) | 2.05 | 0.39 | 1.28 – 2.82 | <0.0001**** |
| age | -0.049 | 0.028 | -0.1 – 0.006 | 0.083 |
| Random effects | Variance |  |  |  |
| Nest | 3.1 |  |  |  |
| Residual | 23.41 |  |  |  |

**Fig 3.**
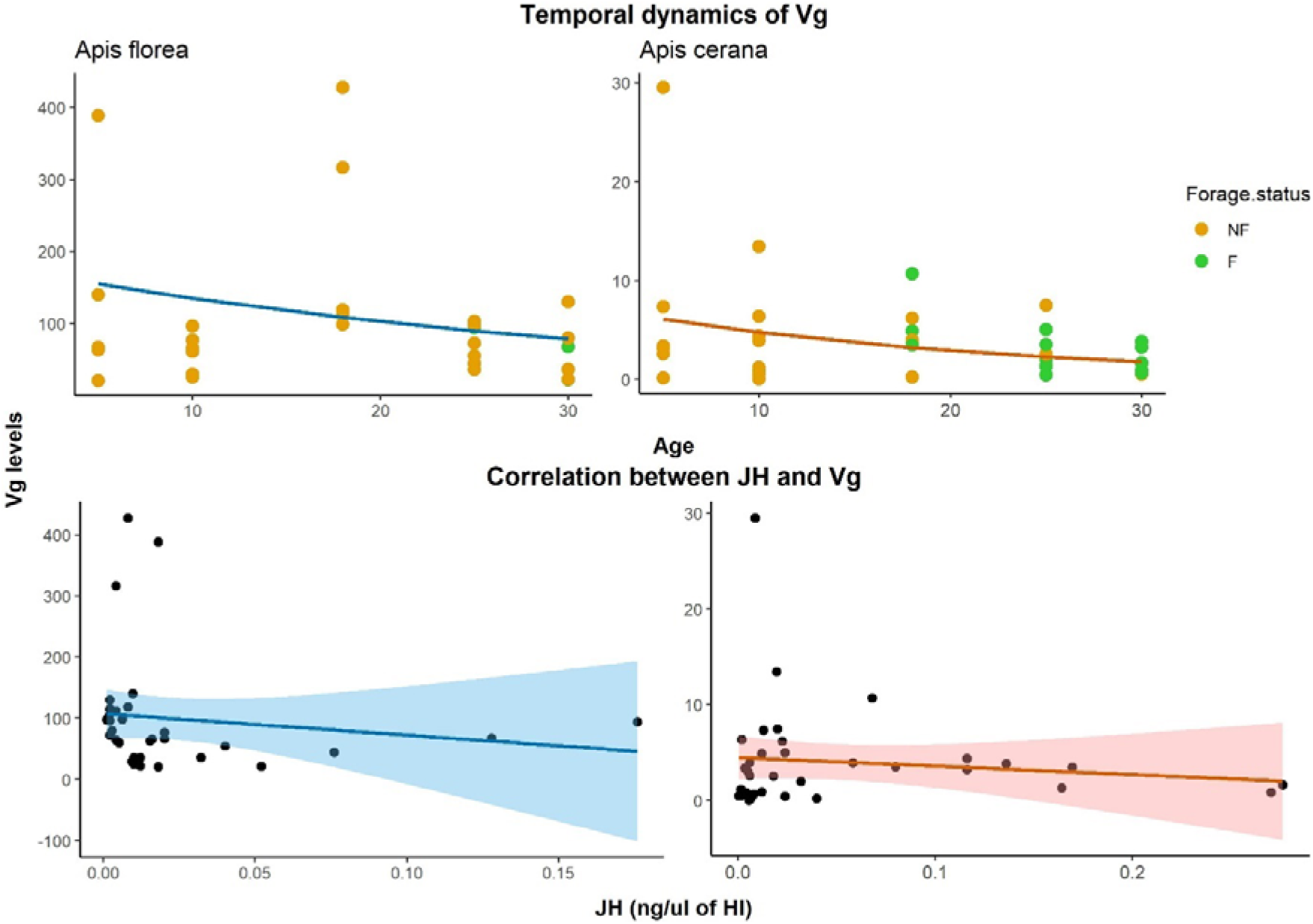
Illustrates the changes in *Vg* titres with age (top panel) and correlation between *Vg* and JH (bottom panel). For the top panel, × axis is the age measured in days and Y axis is the gene expression levels of *Vg* in the bees. Each dot indicates a single bee and orange dots are the bees that had not yet become foragers (NF) by the time of collection while green are the ones that had become (F). Mixed effects models with age as fixed effect, *Vg* levels as response variable and colony as random effect were built. Lines indicate the prediction from the models for the relationship between age and *Vg*. In both species, age did not have a significant effect on *Vg* levels. The bottom panel shows the correlation between JH titres and *Vg* levels and in both species, there was no significant correlation between the two.

Since JH and *Vg* were measured in the same bees, we checked for correlation between JH and *Vg*, but there was no significant correlation in both *A. florea* (coefficient = −0.136, CI = −0.46 - 0.22, P value = 0.449) and *A. cerana* (coefficient = −0.122, CI = −0.43 - 0.22, P value = 0.486) (Fig. 3).

### Hormone titres in nurse bees and foragers

JH titres and *ilp-1* expression levels were significantly higher in foragers than nurses for both

*A. florea* and *A. cerana. Vg* expression levels were not different between nurses and foragers for both species.

#### Juvenile Hormone

Linear mixed effects models (Table 5) showed significant difference between nurses and foragers in their JH titres, with foragers having significantly higher levels than nurses for both *A. florea* and *A. cerana* (Table 5, Fig. 4).

**Table 5:** Table showing the fixed effects, random effects and the estimate with standard error, confidence intervals and P values for the linear mixed effects models to study JH titres in nurses and foragers in *A. florea*. Foragers had significantly higher titre levels of JH than nurses in both *A. florea* and *A. cerana*

| Fixed effects | Estimate | Std. Error | CI | P value |
| --- | --- | --- | --- | --- |
| (Intercept) | 0.042 | 0.026 | -0.025 – 0.11 | 0.17 |
| Social.Role.F | 0.209 | 0.026 | 0.156 – 0.262 | <0.0001**** |
| Random effects | Variance |  |  |  |
| Nest | 0.0011 |  |  |  |
| Residual | 0.0038 |  |  |  |

(b) *A. cerana*
| Fixed effects | Estimate | Std. Error | CI | P value |
| --- | --- | --- | --- | --- |
| (Intercept) | -2.47 | 0.511 | -3.47 – -1.47 | <0.0001**** |
| Social.Role.F | 1.56 | 0.49 | 0.59 – 2.53 | 0.0016** |
| Random effects | Variance |  |  |  |
| Nest | 0.024 |  |  |  |
| Residual | 0.027 |  |  |  |

**Fig 4.**
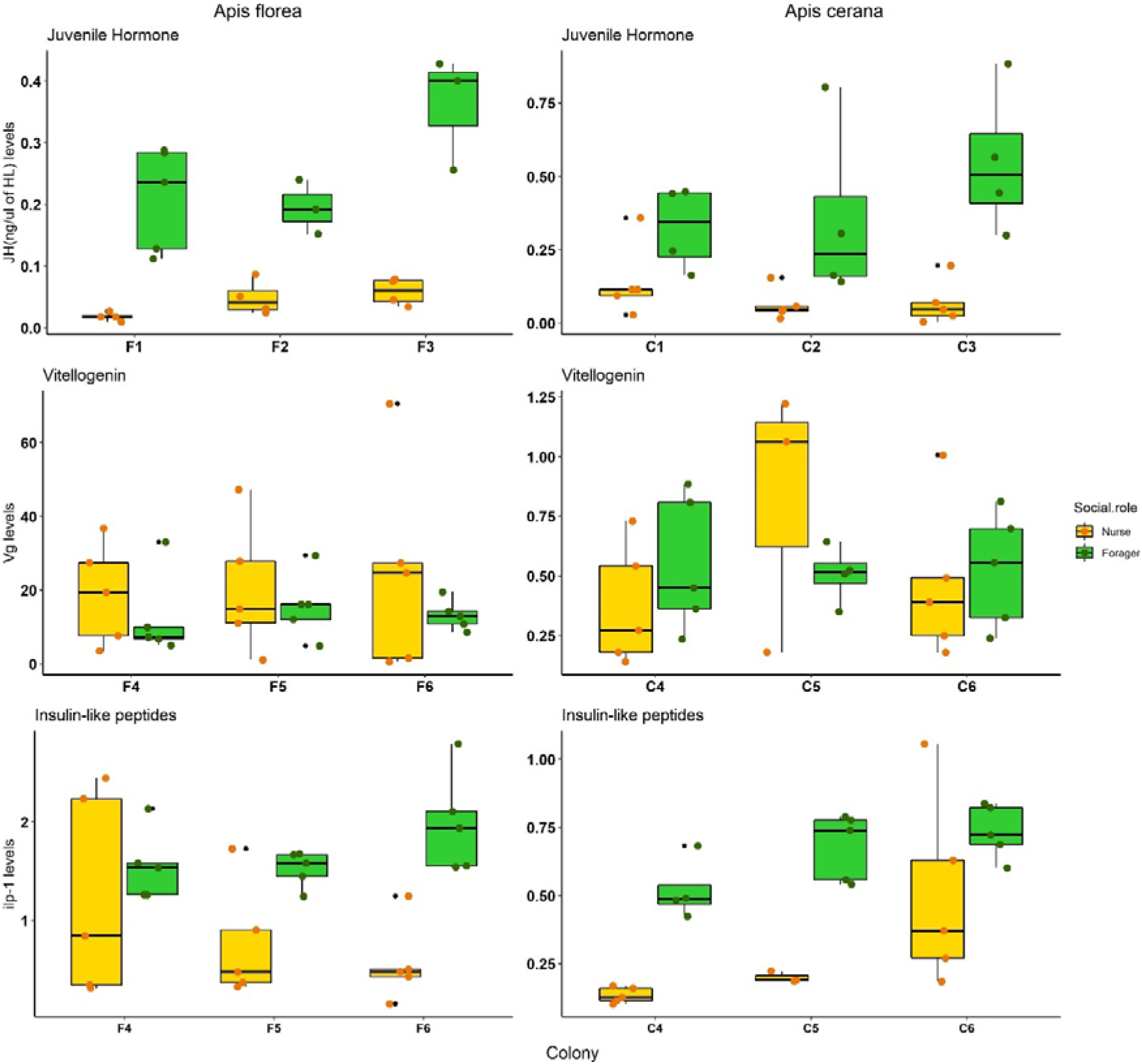
Juvenile hormone titres (top row), vitellogenin expression levels (middle row) and expression levels of insulin-like peptides (bottom row) in nurses and foragers for *A. florea* (left column) and *A. cerana* (right column). Results for colonies are shown separately. Each dot indicates a single bee and orange indicates nurses and green foragers. Regression models were built separately for each of the response variable (JH titres, *Vg* and *ilp-1* expression levels) and for each of the species. The predictor variable was a categorical variable and was the behavioural state, nurse and forager. Although values were scaled for analysis for *Vg* and *ilp-1*, actual values are plotted in the figure. JH and *ilp-1* were significantly higher in foragers than nurses for both species. *Vg* did not differ between nurses and foragers in both the species.

#### Vitellogenin

Linear regression models (Table 6) showed no significant difference in the levels of vitellogenin between nurses and foragers for both *A. florea* and *A. cerana* (Table 6, Fig. 4).

**Table 6:** Table showing the predictor variable and the estimate with standard error, confidence intervals and P values for linear regression models to study *Vg* expression levels in nurses and foragers in *A.florea*. There was no significant difference between foragers and nurses in both *A. florea* and *A. cerana*.

| Predictor | Estimate | Std. Error | CI | P value |
| --- | --- | --- | --- | --- |
| (Intercept) | 0.243 | 0.24 | -0.26 – 0.74 | 0.33 |
| Social.Role.F | -0.486 | 0.35 | -1.19 – 0.22 | 0.17 |
| **R<sup>2</sup>** | **F-statistic** |  |  |  |
| 0.065 | 1.96 |  |  |  |

(b) *A. cerana*
| Predictor | Estimate | Std. Error | CI | P value |
| --- | --- | --- | --- | --- |
| (Intercept) | -0.058 | 0.27 | -0.62 – 0.5 | 0.834 |
| Social.Role.F | 0.111 | 0.38 | -0.66 – 0.89 | 0.77 |
| **R<sup>2</sup>** | **F-statistic** |  |  |  |
| 0.0035 | 0.087 |  |  |  |

#### Insulin-like peptide

Linear regression models showed *ilp-1* was significantly higher in foragers than nurses for both the species of honeybee (Table 7, Fig. 4).

**Table 7:** Table showing the predictor variable and the estimate with standard error, confidence intervals and P values for linear regression models to study *Ilp-1-11* expression levels in nurses and foragers in *A. florea*. *Ilp-1-11* levels were significantly higher in foragers than nurses in both *A. florea* and *A. cerana*.

| Predictor | Estimate | Std. Error | CI | P value |
| --- | --- | --- | --- | --- |
| (Intercept) | -0.56 | 0.20 | -0.98 – -0.15 | 0.0096** |
| Social.Role.F | 1.13 | 0.29 | 0.54 – 1.72 | 0.0005*** |
| **R<sup>2</sup>** | **F-statistic** |  |  |  |
| 0.356 | 15.47 |  |  |  |

(b) *A. cerana*
| Predictor | Estimate | Std. Error | CI | P value |
| --- | --- | --- | --- | --- |
| (Intercept) | -0.58 | 0.22 |  | 0.014* |
| Social.Role.F | 1.09 | 0.30 |  | 0.0013** |
| **R<sup>2</sup>** | **F-statistic** |  |  |  |
| 0.33 | 12.9 |  |  |  |

### Expression levels of foraging related transcription factors

All the TFs measured, i.e., *usp*, *egr-1* and *BR-C*, except for *nautilus* were significantly higher expressed in brains of foragers than nurses for both the species, *A. florea* (Table 8a, Fig. 5) and *A. cerana* (Table 8b, Fig. 5). *Nautilus* was not significantly different between nurses and foragers in *A. florea* and *A. cerana* (Table 8a and b, Fig. 5).

**Fig 5.**
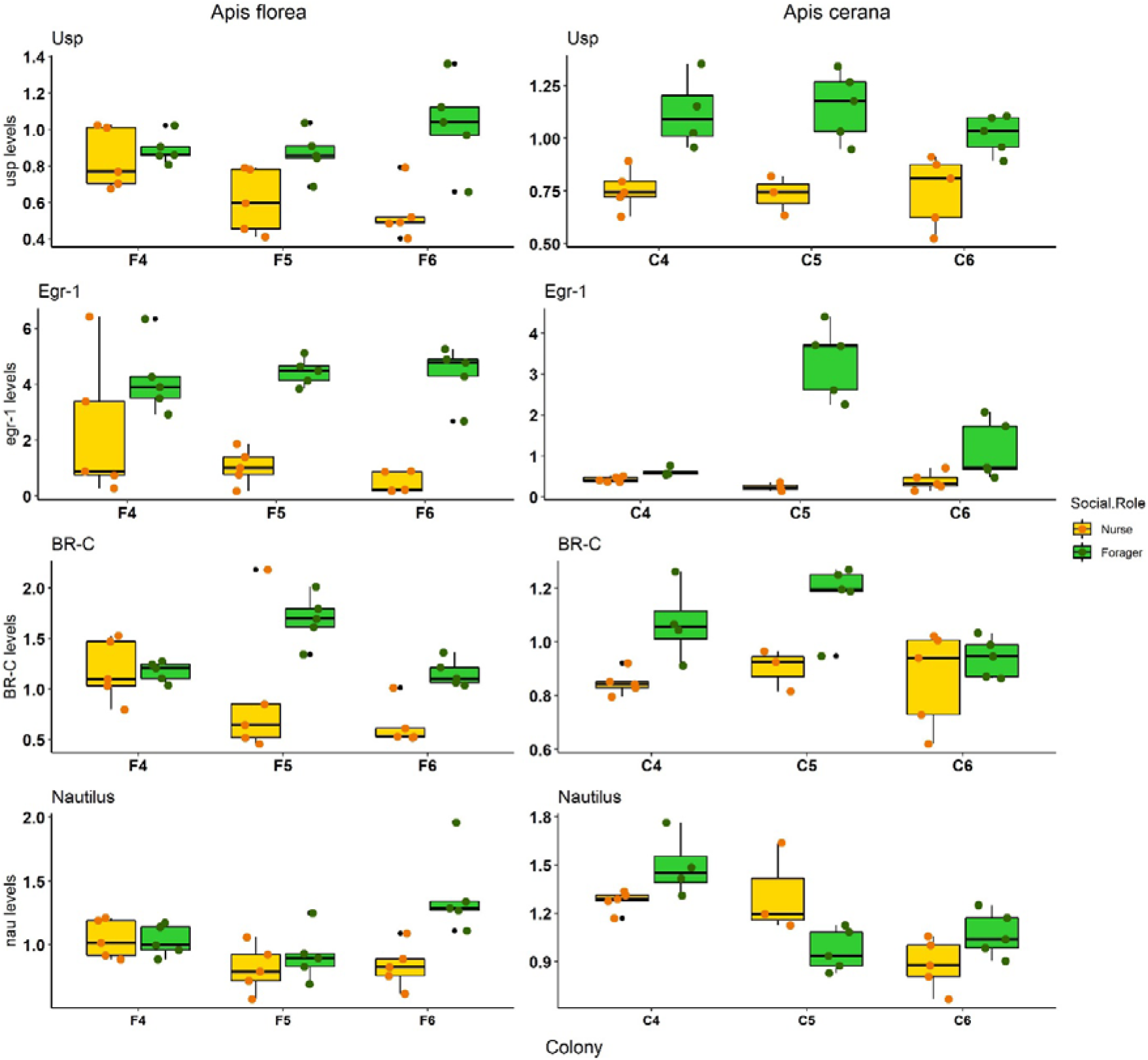
Expression levels for 4 transcription factors (*usp*, *egr-1*, *BR-C and nautilus*) for three colonies in *A. florea* are shown on the left panel and for three colonies for *A. cerana* in the right panel. Dots indicate individual bee values for the TFs. Linear regression models were built for each TFs as the response variable and behavioural state (nurse and forager) as predictor variable. Although values were scaled for analysis, actual values are plotted in the figure. All TFs except *nautilus* were significantly higher in foragers than nurses for both species. *Nautilus* did not differ between nurses and foragers in both *A. florea* and *A. cerana*.

**Table 8a:** Table showing the predictor variable and the estimate with standard error, confidence intervals, R^2^, F-statistic and P values for linear regression models to study 4 TFs, *usp, Egr −1*, *BR-C* and *nautilus* expression levels in nurses and foragers in *A. florea*. All TFs except for *nautilus* were higher in foragers than nurses. *Nautilus* wasn’t different between nurses and foragers.

| Predictor | Estimate | Std. Error | CI | P value |
| --- | --- | --- | --- | --- |
| (Intercept) | -0.54 | 0.21 | -0.97 – -0.11 | 0.015* |
| Social.Role.F | 1.08 | 0.29 | 0.48 – 1.68 | 0.001** |
| $R^2$ | **F-statistic** | | | |
| 0.32 | 13.4 |  |  |  |

(ii) *Egr - 1*
| Predictor | Estimate | Std. Error | CI | P value |
| --- | --- | --- | --- | --- |
| (Intercept) | -0.74 | 0.16 | -1.07 – -0.42 | <0.0001**** |
| Social.Role.F | 1.49 | 0.22 | 1.03 – 1. | <0.0001**** |
| $R^2$ | **F-statistic** | | | |
| 0.61 | 44.7 |  |  |  |

(iii) *BR-C*
| Predictor | Estimate | Std. Error | CI | P value |
| --- | --- | --- | --- | --- |
| (Intercept) | -0.46 | 0.22 | -0.91 – -0.003 | 0.048* |
| Social.Role.F | 0.92 | 0.31 | 0.27 – 1.56 | 0.007** |
| $R^2$ | **F-statistic** | | | |
| 0.23 | 8.5 |  |  |  |

(iv) *Nautilus*
| Predictor | Estimate | Std. Error | CI | P value |
| --- | --- | --- | --- | --- |
| (Intercept) | -0.32 | 0.24 | -0.81 – 0.17 | 0.19 |
| Social.Role.F | 0.64 | 0.34 | -0.06 – 1.33 | 0.07 |
| $R^2$ | **F-statistic** | | | |
| 0.11 | 3.5 |  |  |  |

**Table 8b:**
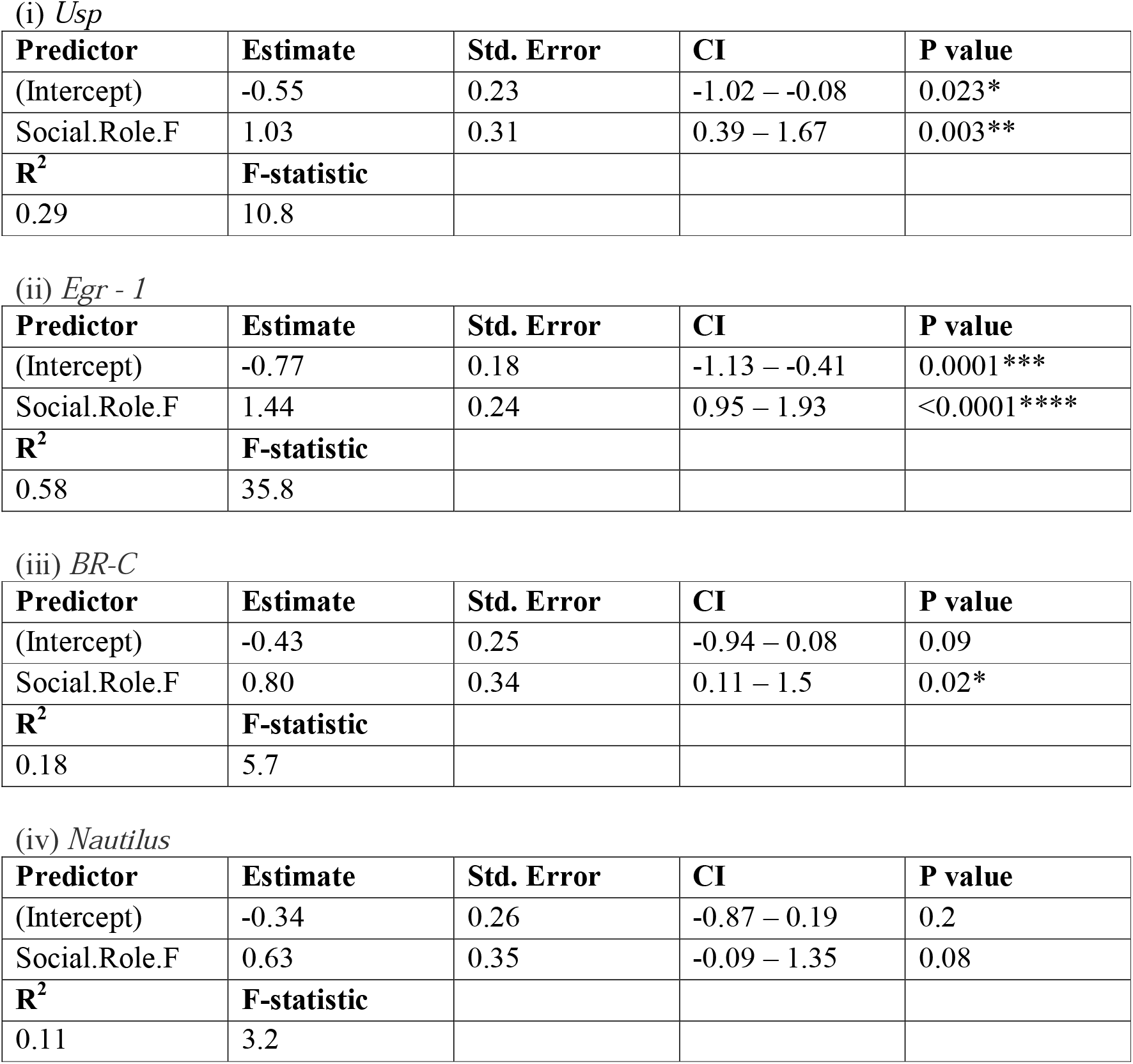
Table showing the predictor variable and the estimate with standard error, confidence intervals, R^2^, F-statistic and P values for linear regression models to study 4 TFs, *usp, Egr −1*, *BR-C* and *nautilus* expression levels in nurses and foragers in *A. florea*. All TFs except for *nautilus* were higher in foragers than nurses. *Nautilus* wasn’t different between nurses and foragers.

## Discussion

Our study provides two significant findings; first, in contrast to earlier studies [46,47], our behavioural observations clearly show that the age of the very first onset of foraging is not different between workers of *A. florea* and *A. cerana*, but the open- and cavity-nesting species differ in the variation of developmental pace among the workers in a colony. Second, all major hormonal and molecular changes which have been associated with the nurse-forager transition are similar between *A. florea* and *A. cerana* and resemble those in *A. mellifera* [14] with a few exceptions. Together, these findings strongly suggest that the regulatory mechanisms underlying behavioural maturation are conserved among honey bees.

As open-nesting species are phylogenetically ancestral, accelerated behavioural maturation has been linked to changes associated with cavity-nesting [47]. One idea is that cavity-nesting allowed to recruit the so-called “curtain bees” for nursing and foraging which enabled an increased worker production and a faster onset of foraging [47]. This hypothesis is supported by the finding that the brood to worker ratio is lower in open-nesting compared to cavity-nesting species [46,50]. In this scenario, open-nesting species are apparently constrained in worker production as they need to keep a large worker force to maintain the curtain [46]. However, studies in our own lab showed that the curtain bees do not constantly stay in the curtain and thus are not necessarily omitted from performing other tasks [49]. The second hypothesis proposed is based on the observation that the brood to worker ratio is higher in cavity- than open-nesting species indicating a higher worker production rate [46,50]. It was argued that this difference affects the capability to replace workers and in particular foragers, which are exposed to higher mortality risks [68–71]. Thus, cavity-nesting allows a faster replacement and consequently also a faster development at the level of the worker population. In contrast open-nesting constrains worker replacement and the more distributed onset of foraging observed in this study might be a strategy to generate a buffer against detrimental losses in the forager population [47,72]. Finally, it is important to note, that workers of temperate populations of *A. mellifera* show a seasonal change in the pace of development. The short-lived “summer bees” become foragers after three weeks, whereas the long-lived “winter bees” show a decelerated behavioural maturation and onset of foraging can be delayed for up to several months [10,73,74]. Whether the winter bees are an adaptation to living in a temperate environment after *A. mellifera* dispersed into Europe from Africa or whether they represent the ancestral character state is an open question [75].

There is ample evidence in *A. mellifera*, that juvenile hormone is certainly one of the major drivers of the pace of behavioural development and time of onset of foraging. JH titres increase with age and are correlated with nursing and foraging behaviour [11,12,14,15,18]. Similarly, we found a significant correlation between age and JH titres for *A. cerana* as well as for *A. florea* workers (Fig. 2). However, the analysis is likely confounded by the behavioural state of the studied workers, i.e., whether they were hive bees or foragers at least in the case of *A. cerana* (see Fig. 2). This finding suggests that the task or the developmental decision to become foragers could positively influence JH titres. In contrast to JH, we did not find a significant effect of age on *Vg* levels in *A. florea* and *A. cerana* (Fig. 3). Consequently, we also did not find a significant correlation between JH and *Vg* in both species (Fig. 3), although there are reports that indicate the inverse relationship between *Vg* and JH in *A. mellifera* [12,28,29,34].

The pattern for JH titres, as well as *Vg* and *ilp-1* expression levels in behaviourally identified nurse bees and foragers in *A. florea* and *A. cerana* are similar and also mainly resemble those in *A. mellifera* [11,12,25]. JH titres and *ilp-1* expression levels were significantly higher in foragers compared to nurses. Our JH results also corroborate a previous study on *A. cerana* which showed higher levels of JH in foragers compared to nurses [45]. In contrast, we did not find differences in the *Vg* expression between nurses and foragers in both species. An earlier study in *A. cerana* had shown that vitellogenin was highly expressed in nurses and then dropped in foragers [40] similar to results in *A. mellifera* [27,75,76]. On the one hand, *Vg* expression levels in workers might be dependent on colony status like presence and amount of brood [77], unfortunately we haven’t checked differences in open brood for the colonies in our study. In addition, there is some evidence in *A. mellifera* that *Vg* titres and interaction between *Vg* and JH are genotype dependent [33]. Finally, to our surprise RNA expression levels of *Vg* in *A. florea* workers were ten to thirty times higher than that in *A. cerana*. In *A. dorsata*, another open-nesting Asian honeybee species, it was found that workers possess on average 20 ovarioles, compared to 3-5 ovarioles in the case of *A. mellifera* and 5-8 in *A. cerana* [78]. These findings may suggest that workers of open-nesting species have a higher reproductive capability and consequently a higher capacity to synthesize and store vitellogenin.

In addition to measuring major hormones involved in behavioural maturation, we also examined whether major brain transcription factors associated with nursing and foraging behaviour in *A. mellifera* showed respective changes in expression between nurses and foragers in *A. florea* and *A. cerana*. With respect to recent publications [13,36,37], we selected two foraging-related transcription factors, ultraspiracle (*usp*) and early growth response 1 (*egr-1*), and two nursing-related transcription factors, broad-complex (*BR-C*) and *nautilus*. *Usp* and *egr-1* both showed robust higher expression levels in foragers than nurse bees in *A. florea* and *A. cerana* similar to what has been found in *A. mellifera* [36–38]. Unexpectedly, expression levels of *BR-C* and *nautilus* in nurse bees and foragers did not show a clear pattern similar to what was observed in *A. mellifera*. *BR-C* was significantly higher in foragers compared to nurses and *nautilus* levels did not significantly differ between nurses and foragers. Interestingly, although the patterns for *BR-C* and *nautilus* did not follow what was observed in *A. mellifera*, they were similar between *A. florea* and *A. cerana* with both species showing the same pattern for both *BR-C* and *nautilus*.

To summarize, all the examined hormonal and molecular changes associated with worker behavioural maturation showed similar characteristics in *A. florea* and *A. cerana*. The only difference appears to be the pace of behavioural development and likely the pace of change in the JH titre. As mentioned above the behavioural studies indicate that the earliest onset of foraging is not different between the two species. What is different is the variation of onset of foraging among the worker population, with *A. cerana* workers showing a narrower distribution in onset of foraging compared to *A. florea*. An accelerated increase in JH titre might be sufficient for such a change, but a lowered JH response threshold could also play a role.

The strong similarities in major hormonal and brain molecular traits among the workers, i.e., nurse bees as well as foragers, of two distantly related honey bee species support the idea that species differences among honey bees are not dramatic but likely subtle. This might be an advantage for comparative studies aiming to identify correlations between behavioural and brain differences. One could start with comparative behavioural experiments but our findings also suggest that the reverse would also work out, which might be more useful for studies on open-nesting bees considering the curtain. Carefully designed comparative molecular studies, e.g., comparing gene regulatory networks or chip-seq experiments, might identify subtle molecular changes suggesting specific behavioural differences.

## Supporting information

Contains supplementary Table 1, figures, S1, S2 and S3

## Author contributions

S.U. participated in the design of the study, conducted the behavioural experiments and mass spectrometry, participated in video analysis and sample collection, performed data analysis and participated in writing the manuscript. A.Sh. designed the primers, participated in RNA extractions, performed the qPCRs and participated in writing the manuscript. D.B. participated in RNA extractions and critically revised the manuscript. A.Su. participated in video analysis, sample collection and critically revised the manuscript. A.B. conceived the study, participated in the design of the study, coordinated the study and participated in writing the manuscript.

## Acknowledgements

We are thankful to Dr. Divya Ramesh and Ccamp mass spec facility for help in mass spectrometry. We would like to thank Dr. Ebi George for help with the statistical analysis. We would also like to thank Dr. Aswathy Nair and Manal Shakeel for helpful comments on the manuscript.

## References

1. Beshers SN, Fewell JH. 2001 Models of Division of Labor in Social Insects. Annual Review of Entomology 46, 413–440. (doi:10.1146/annurev.ento.46.1.413)

2. Oster GF, Wilson EO. 1978 Caste and Ecology in the Social Insects. Princeton University Press.

3. Seeley TD. 1982 Adaptive significance of the age polyethism schedule in honeybee colonies. Behav Ecol Sociobiol 11, 287–293. (doi:10.1007/BF00299306)

4. Wilson EO. 1985 The Sociogenesis of Insect Colonies. Science 228, 1489–1495. (doi:10.1126/science.228.4707.1489)

5. Noirot C, Bordereau C. 1991 Termite polymorphism and and morphogenetic hormones. In Morphogenetic hormones of arthropods, pp. 295–324. New Brunswick: Rutgers University Press.

6. Barredo CG, Gil-Marti B, Deveci D, Romero NM, Martin FA. 2020 Timing the Juvenile-Adult Neurohormonal Transition: Functions and Evolution. Front Endocrinol (Lausanne) 11, 602285. (doi:10.3389/fendo.2020.602285)

7. Fahrbach SE, Robinson GE. 1996 Juvenile Hormone, Behavioral Maturation, and Brain Structure in the Honey Bee. DNE 18, 102–114. (doi:10.1159/000111474)

8. Whitfield CW, Ben-Shahar Y, Brillet C, Leoncini I, Crauser D, LeConte Y, Rodriguez-Zas S, Robinson GE. 2006 Genomic dissection of behavioral maturation in the honey bee. Proc Natl Acad Sci U S A 103, 16068–16075. (doi:10.1073/pnas.0606909103)

9. Johnson BR. 2010 Division of labor in honeybees: form, function, and proximate mechanisms. Behav Ecol Sociobiol 64, 305–316. (doi:10.1007/s00265-009-0874-7)

10. Winston ML. 1987 The biology of the honeybee. Harvard University Press, Cambridge, MA.

11. Robinson GE. 1992 Regulation of Division of Labor in Insect Societies. Annual Review of Entomology 37, 637–665. (doi:10.1146/annurev.en.37.010192.003225)

12. Bloch G, Shpigler H, Wheeler DE, Robinson GE. 2002 Endocrine influences on the organisation of Insect societies. In Hormones, Brain and Behaviour, pp. 1027–1068. San Diego: Academic Press.

13. Hamilton AR, Traniello IM, Ray AM, Caldwell AS, Wickline SA, Robinson GE. 2019 Division of labor in honey bees is associated with transcriptional regulatory plasticity in the brain. Journal of Experimental Biology 222. (doi:10.1242/jeb.200196)

14. Hamilton AR, Shpigler H, Bloch G, Wheeler DE, Robinson GE. 2017 2.19 - Endocrine Influences on Insect Societies. In Hormones, Brain and Behavior (Third Edition) (eds DW Pfaff, M Joëls), pp. 421–451. Oxford: Academic Press. (doi:10.1016/B978-0-12-803592-4.00037-7)

15. Rutz W, Gerig L, Wille H, Lüscher M. 1976 The function of juvenile hormone in adult worker honeybees, Apis mellifera. Journal of Insect Physiology 22, 1485–1491. (doi:10.1016/0022-1910(76)90214-6)

16. Robinson GE. 1987 Regulation of honey bee age polyethism by juvenile hormone. Behav Ecol Sociobiol 20, 329–338. (doi:10.1007/BF00300679)

17. Fahrbach S. 1997 Regulation of age polyethsim in bees and wasps by juvenile hormone. In Advances in the Study of Behavior, pp. 285–316. Academic Press.

18. M. Elekonich M, Schulz DJ, Bloch G, Robinson GE. 2001 Juvenile hormone levels in honey bee (Apis mellifera L.) foragers: foraging experience and diurnal variation. Journal of Insect Physiology 47, 1119–1125. (doi:10.1016/S0022-1910(01)00090-7)

19. Jaycox ER, Skowronek W, Guynn G. 1974 Behavioral Changes in Worker Honey Bees (Apis mellifera) Induced by Injections of a Juvenile Hormone Mimic1. Annals of the Entomological Society of America 67, 529–534. (doi:10.1093/aesa/67.4.529)

20. Jaycox ER. 1976 Behavioral Changes in Worker Honey Bees (Apis mellifera L.) after Injection with Synthetic Juvenile Hormone (Hymenoptera: Apidae). Journal of the Kansas Entomological Society 49, 165–170.

21. Robinson GE. 1985 Effects of a juvenile hormone analogue on honey bee foraging behaviour and alarm pheromone production. Journal of Insect Physiology 31, 277–282. (doi:10.1016/0022-1910(85)90003-4)

22. Sasagawa H, Sasaki M, Okada I. 1989 Hormonal Control of the Division of Labor in Adult Honeybees (Apis mellifera L.)D: I. Effect of Methoprene on Corpora Allata and Hypopharyngeal Gland, and Its α-Glucosidase Activity. Applied Entomology and Zoology 24, 66–77. (doi:10.1303/aez.24.66)

23. Sullivan JP, Fahrbach SE, Harrison JF, Capaldi EA, Fewell JH, Robinson GE. 2003 Juvenile hormone and division of labor in honey bee colonies: effects of allatectomy on flight behavior and metabolism. Journal of Experimental Biology 206, 2287–2296. (doi:10.1242/jeb.00432)

24. Rubenstein DR, Alcock J. 2019 Animal Behaviour. International eleventh edition/Dustin R. Rubenstein, John Alcock. New York. Oxford University Press.

25. Ament SA, Corona M, Pollock HS, Robinson GE. 2008 Insulin signaling is involved in the regulation of worker division of labor in honey bee colonies. PNAS 105, 4226–4231. (doi:10.1073/pnas.0800630105)

26. Ament SA, Velarde RA, Kolodkin M, Moyse D, Robinson GE. 2011 Neuropeptide Y-like signaling and nutritionally-mediated gene expression and behavior in the honey bee. Insect Mol Biol 20, 335–345. (doi:10.1111/j.1365-2583.2011.01068.x)

27. Engels W. 1974 Occurrence and Significance of Vitellogenins in Female Castes of Social Hymenoptera. American Zoologist 14, 1229–1237. (doi:10.1093/icb/14.4.1229)

28. Hartfelder K, Engels W. 1998 2 Social Insect Polymorphism: Hormonal Regulation of Plasticity in Development and Reproduction in the Honeybee. In Current Topics in Developmental Biology (eds RA Pedersen, GP Schatten), pp. 45–77. Academic Press. (doi:10.1016/S0070-2153(08)60364-6)

29. Fahrbach SE, Giray T, Robinson GE. 1995 Volume Changes in the Mushroom Bodies of Adult Honey Bee Queens. Neurobiology of Learning and Memory 63, 181–191. (doi:10.1006/nlme.1995.1019)

30. Pinto LZ, Bitondi MMG, Simões ZLP. 2000 Inhibition of vitellogenin synthesis in Apis mellifera workers by a juvenile hormone analogue, pyriproxyfen. Journal of Insect Physiology 46, 153–160. (doi:10.1016/S0022-1910(99)00111-0)

31. Barchuk AR, Bitondi MMG, Simões ZLP. 2002 Effects of juvenile hormone and ecdysone on the timing of vitellogenin appearance in hemolymph of queen and worker pupae of Apis mellifera. J Insect Sci 2.

32. Corona M, Velarde RA, Remolina S, Moran-Lauter A, Wang Y, Hughes KA, Robinson GE. 2007 Vitellogenin, juvenile hormone, insulin signaling, and queen honey bee longevity. PNAS 104, 7128–7133. (doi:10.1073/pnas.0701909104)

33. Ihle KE, Page RE, Frederick K, Fondrk MK, Amdam GV. 2010 Genotype effect on regulation of behaviour by vitellogenin supports reproductive origin of honeybee foraging bias. Animal Behaviour 79, 1001–1006. (doi:10.1016/j.anbehav.2010.02.009)

34. Amdam GV, Omholt SW. 2003 The hive bee to forager transition in honeybee colonies: the double repressor hypothesis. Journal of Theoretical Biology 223, 451–464. (doi:10.1016/S0022-5193(03)00121-8)

35. Whitfield CW, Cziko A-M, Robinson GE. 2003 Gene Expression Profiles in the Brain Predict Behavior in Individual Honey Bees. Science 302, 296–299. (doi:10.1126/science.1086807)

36. Chandrasekaran S, Ament SA, Eddy JA, Rodriguez-Zas SL, Schatz BR, Price ND, Robinson GE. 2011 Behavior-specific changes in transcriptional modules lead to distinct and predictable neurogenomic states. PNAS 108, 18020–18025. (doi:10.1073/pnas.1114093108)

37. Khamis AM et al. 2015 Insights into the Transcriptional Architecture of Behavioral Plasticity in the Honey Bee Apis mellifera. Scientific Reports 5, 11136. (doi:10.1038/srep11136)

38. Ament SA et al. 2012 The Transcription Factor Ultraspiracle Influences Honey Bee Social Behavior and Behavior-Related Gene Expression. PLOS Genetics 8, e1002596. (doi:10.1371/journal.pgen.1002596)

39. Shah A, Jain R, Brockmann A. 2018 Egr-1: A Candidate Transcription Factor Involved in Molecular Processes Underlying Time-Memory. Front. Psychol. 9. (doi:10.3389/fpsyg.2018.00865)

40. Diao Q et al. 2018 Genomic and transcriptomic analysis of the Asian honeybee Apis cerana provides novel insights into honeybee biology. Scientific Reports 8, 822. (doi:10.1038/s41598-017-17338-6)

41. Oppenheim S et al. 2020 Whole Genome Sequencing and Assembly of the Asian Honey Bee Apis dorsata. Genome Biology and Evolution 12, 3677–3683. (doi:10.1093/gbe/evz277)

42. Wang AR, Kim MJ, Park JS, Choi YS, Thapa R, Lee KY, Kim I. 2013 Complete mitochondrial genome of the dwarf honeybee, Apis florea (Hymenoptera: Apidae). Mitochondrial DNA 24, 208–210. (doi:10.3109/19401736.2012.744986)

43. Jourjine N, Hoekstra HE. 2021 Expanding evolutionary neuroscience: insights from comparing variation in behavior. Neuron (doi:10.1016/j.neuron.2021.02.002)

44. Sen Sarma M, Whitfield CW, Robinson GE. 2007 Species differences in brain gene expression profiles associated with adult behavioral maturation in honey bees. BMC Genomics 8, 202. (doi:10.1186/1471-2164-8-202)

45. Huang Z, Kuang H, Kuang B, Qin Y. 2001 Juvenile hormone and division of labour in Apis cerana. Internal. Bec. Rec. Assoc. Cardiff, UK.

46. Seeley TD, Seeley RH, Akratanakul P. 1982 Colony Defense Strategies of the Honeybees in Thailand. Ecological Monographs 52, 43–63. (doi:10.2307/2937344)

47. Dyer FC, Seeley TD. 1991 Nesting behavior and the evolution of worker tempo in four honey bee species. (Ecology)., ECOLOGY 72, 156–170.

48. Oldroyd BP, Wongsiri S. 2006 Asian honey bees biology, conservation, and human interactions. Cambridge, MA: Harvard University Press.

49. Bhagavan H, Muthmann O, Brockmann A. 2016 Structural and temporal dynamics of the bee curtain in the open-nesting honey bee species, Apis florea. Apidologie 47, 749–758. (doi:10.1007/s13592-016-0428-8)

50. Seeley TD, Morse RA. 1976 The nest of the honey bee (Apis mellifera L.). Ins. Soc 23, 495–512. (doi:10.1007/BF02223477)

51. Hepburn HR, Pirk CWW, Duangphakdee O. 2014 Honeybee Nests: Composition, Structure, Function. Berlin Heidelberg: Springer-Verlag. (doi:10.1007/978-3-642-54328-9)

52. Seeley TD, Visscher PK. 1985 Survival of honeybees in cold climates: the critical timing of colony growth and reproduction. Ecological Entomology 10, 81–88. (doi:https://doi.org/10.1111/j.1365-2311.1985.tb00537.x)

53. Seeley TD. 1985 Honeybee Ecology. Princeton University Press.

54. Bhagavan H, Brockmann A. 2019 Apis florea workers show a prolonged period of nursing behavior. Apidologie 50, 63–70. (doi:10.1007/s13592-018-0618-7)

55. Brouwers EVM, Ebert R, Beetsma J. 1987 Behavioural and Physiological Aspects of Nurse Bees in Relation to the Composition of Larval Food During Caste Differentiation in the Honeybee. Journal of Apicultural Research 26, 11–23. (doi:10.1080/00218839.1987.11100729)

56. Huang Z-Y, Otis GW. 1991 Nonrandom visitation of brood cells by worker honey bees (Hymenoptera: Apidae). J Insect Behav 4, 177–184. (doi:10.1007/BF01054610)

57. Johnson BR. 2008 Within-nest temporal polyethism in the honey bee. Behav Ecol Sociobiol 62, 777–784. (doi:10.1007/s00265-007-0503-2)

58. Ares AM, Nozal MJ, Bernal JL, Martín-Hernández R, M. Higes, Bernal J. 2012 Liquid chromatography coupled to ion trap-tandem mass spectrometry to evaluate juvenile hormone III levels in bee hemolymph from Nosema spp. infected colonies. Journal of Chromatography B 899, 146–153. (doi:10.1016/j.jchromb.2012.05.016)

59. Scholl C, Wang Y, Krischke M, Mueller MJ, Amdam GV, Rössler W. 2014 Light exposure leads to reorganization of microglomeruli in the mushroom bodies and influences juvenile hormone levels in the honeybee. Dev Neurobiol 74, 1141–1153. (doi:10.1002/dneu.22195)

60. Singh AS, Shah A, Brockmann A. 2017 Honey bee foraging induces upregulation of early growth response protein 1, hormone receptor 38 and candidate downstream genes of the ecdysteroid signalling pathway. Insect Mol Biol, n/a-n/a. (doi:10.1111/imb.12350)

61. R Core Team. 2018 R: A language and environment for statistical computing.

62. RStudio Team. 2016 RStudio: Integrated Development Environment for R.

63. Bates D, Mächler M, Bolker B, Walker S. 2015 Fitting Linear Mixed-Effects Models Using lme4. Journal of Statistical Software 67, 1–48. (doi:10.18637/jss.v067.i01)

64. Rigby R, Stasinopoulos D. 2005 Generalized additive models for location, scale and shape,(with discussion). Applied Statistics 54, 507–554.

65. Venables W, Ripley B. 2002 Modern Applied Statistics with S. Fourth. New York: Springer.

66. Wickham H. 2016 ggplot2: Elegant Graphics for Data Analysis. 2nd edn. Springer International Publishing. (doi:10.1007/978-3-319-24277-4)

67. Hartig F. In press. DHARMa: Residual Diagnostics for Hierarchical (Multi-Level / Mixed) Regression Models.

68. Visscher PK, Dukas R. 1997 Survivorship of foraging honey bees. Insectes soc. 44, 1–5. (doi:10.1007/s000400050017)

69. Neukirch A. 1982 Dependence of the life span of the honeybee (Apis mellifica) upon flight performance and energy consumption. J Comp Physiol B 146, 35–40. (doi:10.1007/BF00688714)

70. Page RE, Peng CY-S. 2001 Aging and development in social insects with emphasis on the honey bee, Apis mellifera L. Experimental Gerontology 36, 695–711. (doi:10.1016/S0531-5565(00)00236-9)

71. Rueppell O, Bachelier C, Fondrk MK, Page RE. 2007 Regulation of life history determines lifespan of worker honey bees (Apis mellifera L.). Experimental Gerontology 42, 1020–1032. (doi:10.1016/j.exger.2007.06.002)

72. Young AM, Brockmann A, Dyer FC. 2021 Adaptive tuning of the exploitation-exploration trade-off in four honey bee species. Behav Ecol Sociobiol 75, 20. (doi:10.1007/s00265-020-02938-6)

73. Döke MA, Frazier M, Grozinger CM. 2015 Overwintering honey bees: biology and management. Current opinion in insect science 10, 185–193. (doi:10.1016/j.cois.2015.05.014)

74. Seeley TD, Visscher PK. 1985 Survival of honeybees in cold climates: the critical timing of colony growth and reproduction. Ecological Entomology 10, 81–88. (doi:https://doi.org/10.1111/j.1365-2311.1985.tb00537.x)

75. Amdam GV, Norberg K, Omholt SW, Kryger P, Lourenço AP, Bitondi MMG, Simões ZLP. 2005 Higher vitellogenin concentrations in honey bee workers may be an adaptation to life in temperate climates. Insect. Soc. 52, 316–319. (doi:10.1007/s00040-005-0812-2)

76. Amdam GV, Norberg K, Hagen A, Omholt SW. 2003 Social exploitation of vitellogenin. PNAS 100, 1799–1802. (doi:10.1073/pnas.0333979100)

77. Eyer M, Dainat B, Neumann P, Dietemann V. 2017 Social regulation of ageing by young workers in the honey bee, Apis mellifera. Experimental Gerontology 87, 84–91. (doi:10.1016/j.exger.2016.11.006)

78. Velthuis HHW, Clement J, Morse RA, Laigo FM. 1971 The Ovaries of Apis Dorsata Workers and Oueens from the Philippines. Journal of Apicultural Research 10, 63–66. (doi:10.1080/00218839.1971.11099672)

