## Supplementary material for "Conserved hormonal and molecular mechanisms underlying behavioural maturation in open- and cavity-nesting honey bees": Contains supplementary Table 1, figures, S1, S2 and S3

**Table S1. Details of colonies used for experiments**

| **Colony name** | **Species** | **Number of frames** | **Experiment** |
| --- | --- | --- | --- |
| Cerana 1 | *A. cerana* | 4 (observed only 2) | Onset of foraging |
| Cerana 2 | *A. cerana* | 4 (observed only 2) | Onset of foraging |
| Florea 1 | *A. florea* | N/A | Onset of foraging |
| Florea 2 | *A. florea* | N/A | Onset of foraging |
| CJ1 | *A. cerana* | 6 | Temporal dynamics of JH and *Vg* |
| CJ2 | *A. cerana* | 6 | Temporal dynamics of JH and *Vg* |
| FJ1 | *A. florea* | N/A | Temporal dynamics of JH and *Vg* |
| FJ2 | *A. florea* | N/A | Temporal dynamics of JH and *Vg* |
| C1 | *A. cerana* | 5 | Nurse-Forager comparison (JH) |
| C2 | *A. cerana* | 5 | Nurse-Forager comparison (JH) |
| C3 | *A. cerana* | 5 | Nurse-Forager comparison (JH) |
| F1 | *A. florea* | N/A | Nurse-Forager comparison (JH) |
| F2 | *A. florea* | N/A | Nurse-Forager comparison (JH) |
| F3 | *A. florea* | N/A | Nurse-Forager comparison (JH) |
| C4 | *A. cerana* | 5 | Nurse-Forager comparison (*Vg*, *Ilp-1-1*, TFs) |
| C5 | *A. cerana* | 5 | Nurse-Forager comparison (*Vg*, *Ilp-1-1*, TFs) |
| C6 | *A. cerana* | 5 | Nurse-Forager comparison (*Vg*, *Ilp-1-1*, TFs) |
| F4 | *A. florea* | N/A | Nurse-Forager comparison (*Vg*, *Ilp-1-1*, TFs) |
| F5 | *A. florea* | N/A | Nurse-Forager comparison (*Vg*, *Ilp-1-1*, TFs) |
| F6 | *A. florea* | N/A | Nurse-Forager comparison (*Vg*, *Ilp-1-1*, TFs) |

| Day of observation | Number of bees that disappeared |
| --- | --- |
| 1 | 5 |
| 2 | 3 |
| 3 | 0 |
| 4 | 2 |
| 5 | 1 |
| 6 | 0 |
| 7 | 3 |
| 8 | 1 |
| 9 | 0 |
| 10 | 0 |
| 11 | 0 |
| 12 | 0 |
| 13 | 0 |
| 14 | 3 |
| 15 | 0 |
| 16 | 3 |
| 17 | 4 |
| 18 | 0 |
| 19 | 2 |
| 20 | 0 |
| 21 | 0 |

Fig. S1. Individuals that disappeared on each day (the day they were last seen) during observations in the colony, Cerana 1.

| Day of observation | Number of bees that disappeared |
| --- | --- |
| 1 | 17 |
| 2 | 5 |
| 3 | 3 |
| 4 | 2 |
| 5 | 6 |
| 6 | 1 |
| 7 | 0 |
| 8 | 0 |
| 9 | 4 |
| 10 | 3 |
| 11 | 4 |
| 12 | 1 |
| 13 | 1 |
| 14 | 1 |
| 15 | 1 |
| 16 | 3 |
| 17 | 1 |
| 18 | 3 |
| 19 | 0 |
| 20 | 2 |
| 21 | 2 |

Fig. S2. Individuals that disappeared on each day (the day they were last seen) during observations in the colony, Cerana 2.

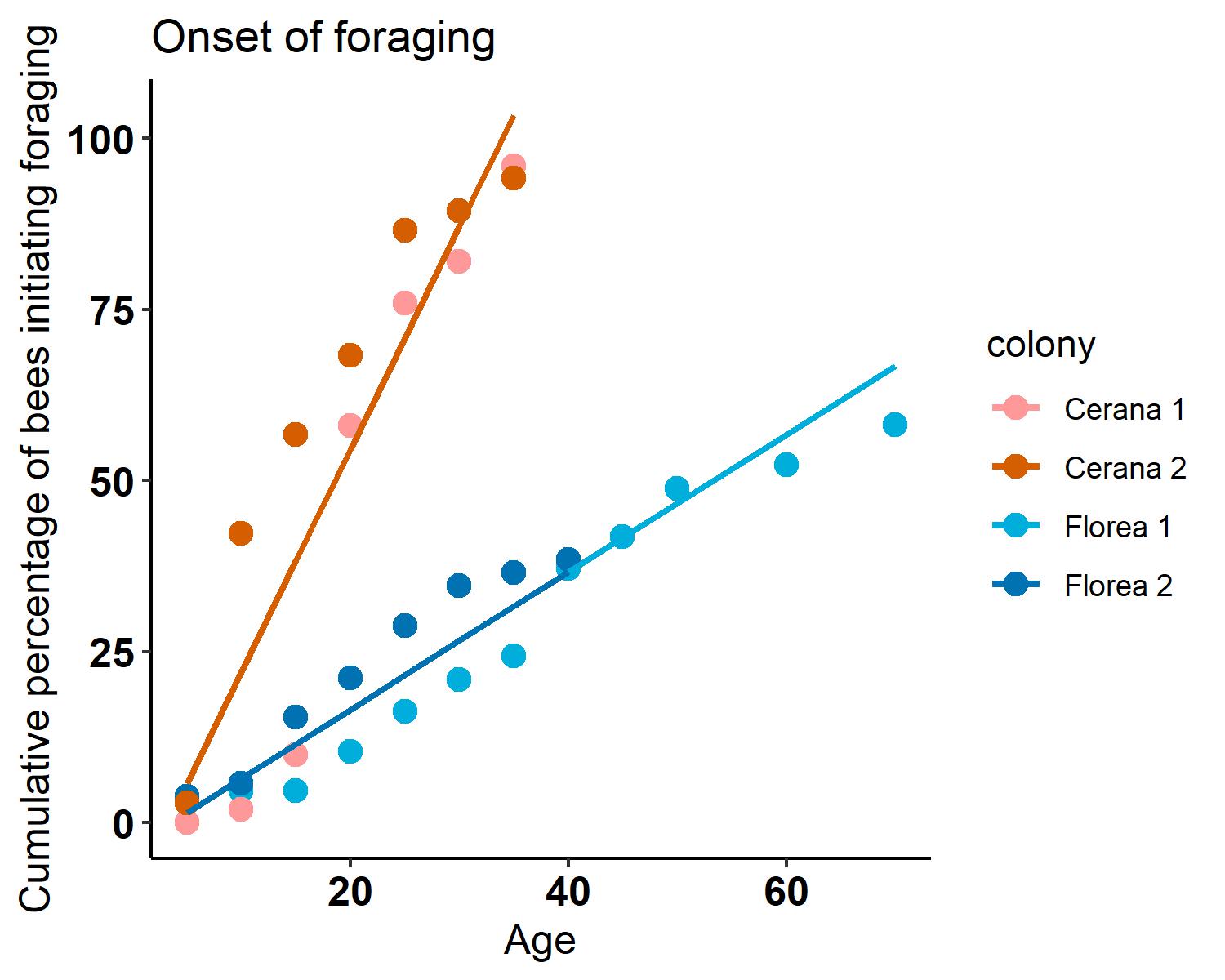

Fig. S3. Cumulative percentage of bees that became foragers out of the number of bees that were still alive (which was 50 bees for Cerana 1 and 104 bees for Cerana 2) by the end of the observations in the case of *A. cerana*. 48 bees out of 50 remaining bees (96%) had become foragers in Cerana 1 and 98 bees out of 104 remaining bees (94%) had become foragers in Cerana 2. In the case of *A. florea* since observations were not possible on the comb the exact age/day of disappearance of bees was difficult to calculate and so cumulative percentage of bees that became foragers out of the total number of bees that were spotted on the curtain (which was 86 for Florea 1 and 52 for Florea 2) during observations was calculated. 50 bees out of 86 bees (58%) spotted on curtain became foragers in Florea 1 and 20 bees out of 52 bees (38%) spotted on curtain became foragers in Florea 2.
